# Ancient ancestry informative markers for identifying fine-scale ancient population structure in Eurasians

**DOI:** 10.1101/333690

**Authors:** Umberto Esposito, Ranajit Das, Mehdi Pirooznia, Eran Elhaik

## Abstract

The rapid accumulation of ancient human genomes from various areas and time periods potentially allows the expansion of studies of biodiversity, biogeography, forensics, population history, and epidemiology into past populations. However, most ancient DNA (aDNA) data were generated through microarrays designed for modern-day populations known to misrepresent the population structure. Past studies addressed these problems using ancestry informative markers (AIMs). However, it is unclear whether AIMs derived from contemporary human genomes can capture ancient population structure and whether AIM finding methods are applicable to ancient DNA (aDNA) provided that the high missingness rates in ancient, oftentimes haploid, DNA can also distort the population structure. Here, we define ancient AIMs (aAIMs) and develop a framework to evaluate established and novel AIM-finding methods in identifying the most informative markers. We show that aAIMs identified by a novel principal component analysis (PCA)-based method outperforms all competing methods in classifying ancient individuals into populations and identifying admixed individuals. In some cases, predictions made using the aAIMs were more accurate than those made with a complete marker set. We discuss the features of the ancient Eurasian population structure and strategies to identify aAIMs. This work informs the design of population microarrays and the interpretation of aDNA results.

## Introduction

### Toward high-resolution population models using ancient samples

Over the past decade, genomic techniques have been reshaping our fundamental understanding of human prehistory and origins [1]. Ancient DNA (aDNA) human genomes have aided in investigations of population structure, human migration, human adaptation, agricultural lifestyle, and disease co-evolution [2]. Ancient genome studies have already accelerated progress in the search for genetic variations underlying the inheritance of adaptations and forensics traits. Recently, Cassidy et al. [3] tested the allelic association of cystic fibrosis and hemochromatosis in ancient Irish samples, expanding genetic epidemiology onto ancient genomes. Such studies can potentially identify new risk factors for rare diseases.

### Next generation sequencing technologies to study ancient DNA

Whole genome sequencing and SNP microarrays are the two leading approaches to aDNA sequencing. Although the former is preferable as it provides more data, by late 2017, only a quarter of the 1,100 sequenced ancient humans were whole genomes. The vast majority of genomes (762) were captured by SNP microarrays [2], mainly the Human Origins [4] and Illumina 610-Quad arrays [5, 6] – neither of which were designed for ancient humans – making it particularly challenging to identify and control for ancient population structure.

Single nucleotide polymorphism (SNP) genotyping microarrays were originally developed to detect phenotype-genotype associations in association-, admixture-, identity by descent-mapping and alike studies. It was not until later that SNP microarrays were employed in population genetic studies aimed at inferring population structure through principal component analyses (PCAs), admixture-like programs, and other tools aimed at predicting group membership. It soon became clear that the allele frequency spectrum obtained through microarrays is more skewed for some populations than for other ones due to the choice of SNP panels [7]. The Human Origins and various Illumina microarrays (including the Illumina Human 660W-Quad, which is very similar to the Illumina 610-Quad array) were shown to distort the population structure for modern day populations compared to larger genomic databases and underreport the biodiversity compared to microarrays customized for population genetics [8, 9], which results in an ascertainment bias.

### The problems of ascertainment bias and population stratification in aDNA

Any inference of identity in archeological studies is fraught with difficulties. Carbon dating requires extracting organic material from fossil bones and authenticating it as composed of degraded proteins – a process highly subjectable to contamination which yields erroneous estimates [10]. The identification of ‘cultures’ from archaeological remains and their association with past population groups is also inadequate [11]. Population genetic studies suffer from similar problems due to ascertainment bias, which can distort measures of human diversity, bias population genetic inferences, and alter the conclusions in unexpected ways [12]. Ascertainment bias is a major concern in genetic, biomedical, and evolutionary studies particularly in the absence of an established population structure model for either modern-day or ancient populations.

The difficulties to establish an acceptable population model are partially due to our incomplete knowledge of human population biodiversity in the past and present. Consequently, modern-day populations are oftentimes assumed to be the parental populations of the modern-day population of interest, which results in population stratification. This problem arises due to differences in the allele frequencies of unknown case\control subpopulations due to separate demographic histories (not biological processes). A misunderstanding of the population structure necessitates mismatched cases and controls, which introduces genetic heterogeneity into the analysis that can lead to spurious associations and obscure the true association [13]. Thereby, the phenotypes of interest (e.g., risk loci or drug response) may differ between these sub-populations and bias the association analyses by generating false positives [14].

These problems have been well known for a long time [15], and statistical remedies have been proposed; however, they were all tailored for modern-day data and did not address the conceptual problems. It is now clear that population models should consider aDNA data and the unique challenges they pose, namely haploidy and high missingness [1].

### The use of ancestry informative markers (AIMs) in genetics

Past studies resolved, to a large extent, the problems faced in aDNA analyses with ancestry informative markers (AIMs). AIMs are SNPs which exhibit large variation in minor allele frequencies (MAF) among populations. Over the past two centuries and to this date, geneticists have scoured genomes for these patterns and produced numerous AIM sets to determine an individual’s ancestry, detect stratification in biomedical studies, infer geographic structure, find risk loci in a candidate region, and localize biogeographical origins [e.g., 8, 9, 16–18]. AIM panels can delineate population structure in a cost effective manner by detecting variation in individual ancestry that can confound methods like Mendelian Randomization trials, association analyses, and forensic investigations in increasing false positive results or reducing power [e.g., 19].

Although initially preferred due to the high cost of sequencing, which has decreased with time, AIMs are still highly used in forensics, carrier screening, and biogeography in both microarrays [e.g., 8, 20] and whole genomic data [21]. Admixture mapping is another powerful method to map phenotypic variation or diseases that show differential risk by ancestry. The mapping takes advantage of higher densities of genetic variants and extensions to admixed populations that exhibit strong differences in prevalence across populations [22]. It is therefore necessary to have a large number of AIMs throughout the genome to allow for inference of local chromosomal ancestry blocks.

Despite their high prevalence, it was never clear which AIMs should be used. All AIM panels have limitations [23], and it is unknown whether established AIMs would be informative for ancient studies. The characteristics of ideal AIMs remain contentious with some authors preferring common SNPs (minor allele frequency >1%) [24], SNPs with high *F*_*ST*_ [25], SNPs with high pairwise MAF between populations [23], or SNPs that satisfy several criteria. Consequently, AIMs do not overlap across studies. Of the 21 AIM datasets, reviewed by Pakstis et al. [26], only 1397 AIMs appeared in at least two sets. Finally, studies typically show that AIMs can separate populations or broadly classify individuals into subcontinental populations rather than showing that AIMs can capture the population structure of the complete SNP set or allow fine-population mapping. Given the uncertainties surrounding AIMs, their potential incompatibility to capture ancient structure and admixtures, and the challenges imposed by aDNA data, it is unclear whether, if at all, AIM-finding methods or AIMs can be utilized to study ancient population structure.

### Ancient ancestry informative markers (aAIMs) to define ancient population structure

Unlike modern-DNA, aDNA allows for the construction of AIM panels from the actual parental populations of modern-day people and can, therefore, refine estimates of population structure. To overcome some of the aforementioned problems with aDNA data, we defined ancient ancestry informative markers (aAIMs) as SNPs that vary in their MAF across ancient populations (Figure 1) and attempted to identify and validate the first autosomal aAIMs to improve the inference of ancient population structure. Since AIM-finding tools were never tested on aDNA, it is necessary to first compare their ability in finding informative markers. For that, we interrogated a comprehensive dataset of 302 ancient genomes grouped into 21 populations from Europe, the Middle East, and North Eurasia. This dataset was used to compare the ability of different methods to identify aAIMs that can best capture the population structure and identify admixed individuals. These methods are: two existing AIM finding algorithms (Infocalc [27] and Wright’s *F*_*ST*_ [28, 29]), three novel Admixture- and PCA-based algorithms and two random SNP sets. First, we derived summary statistics using these AIM candidates. Then, we compared the performances of the best aAIM set and the complete SNP set in classifying individuals to populations and identifying two-way admixed individuals (Figure 2). Our study offers a methodological framework to evaluate AIMs, contrasts different AIM-finding strategies, reports the first set of aAIMs, and demonstrates that in some cases they provide more reliable predictions than the complete SNP set.

**Figure 1.**
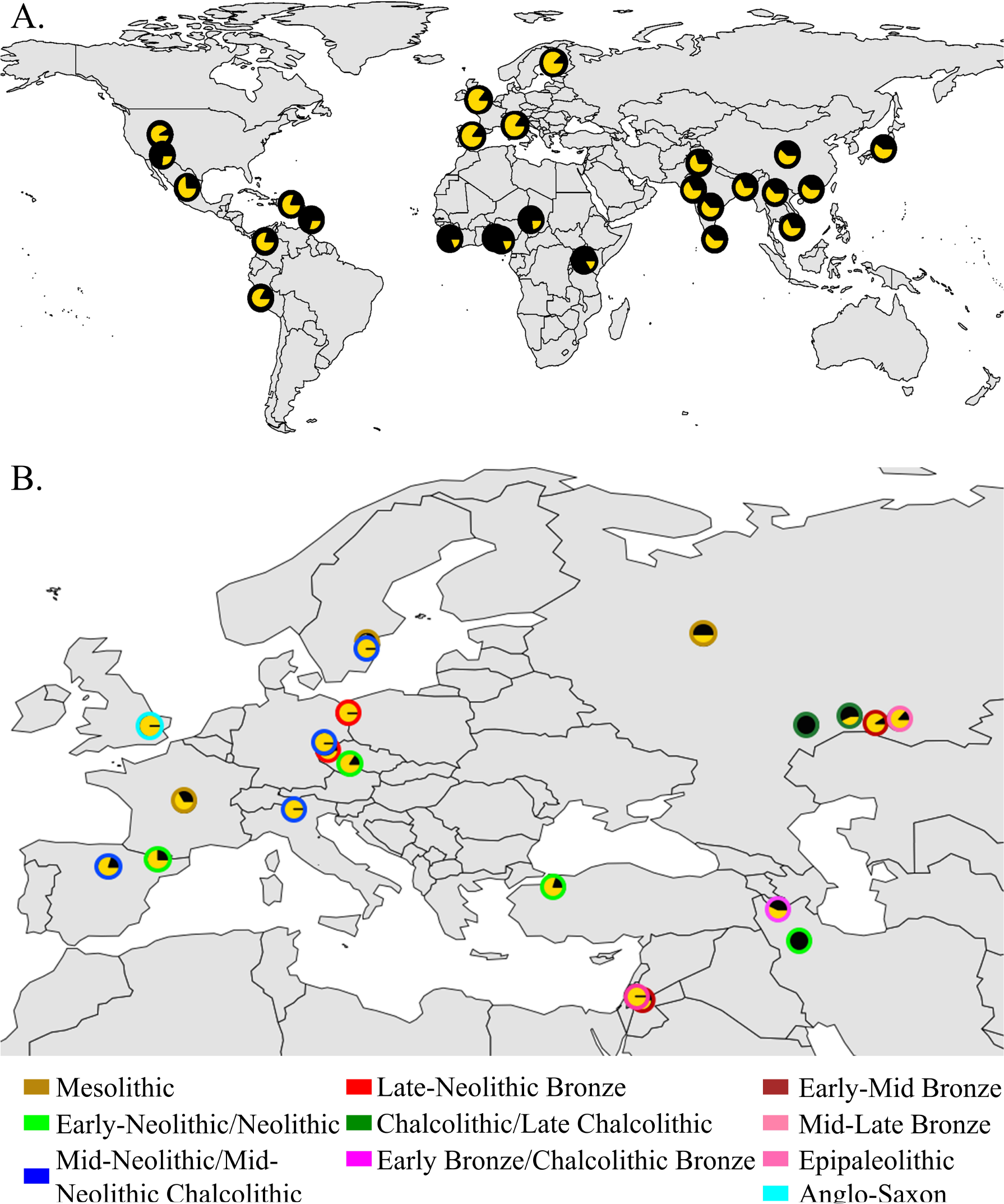
Geographic distribution of the highly differentiated rs7896530 in modern-day (A) and ancient (B) populations. The geographic distributions of the T (black) and G (yellow) alleles were obtained from the Geography of Genetic Variants Browser [50] and Table S1, respectively.

**Figure 2.**
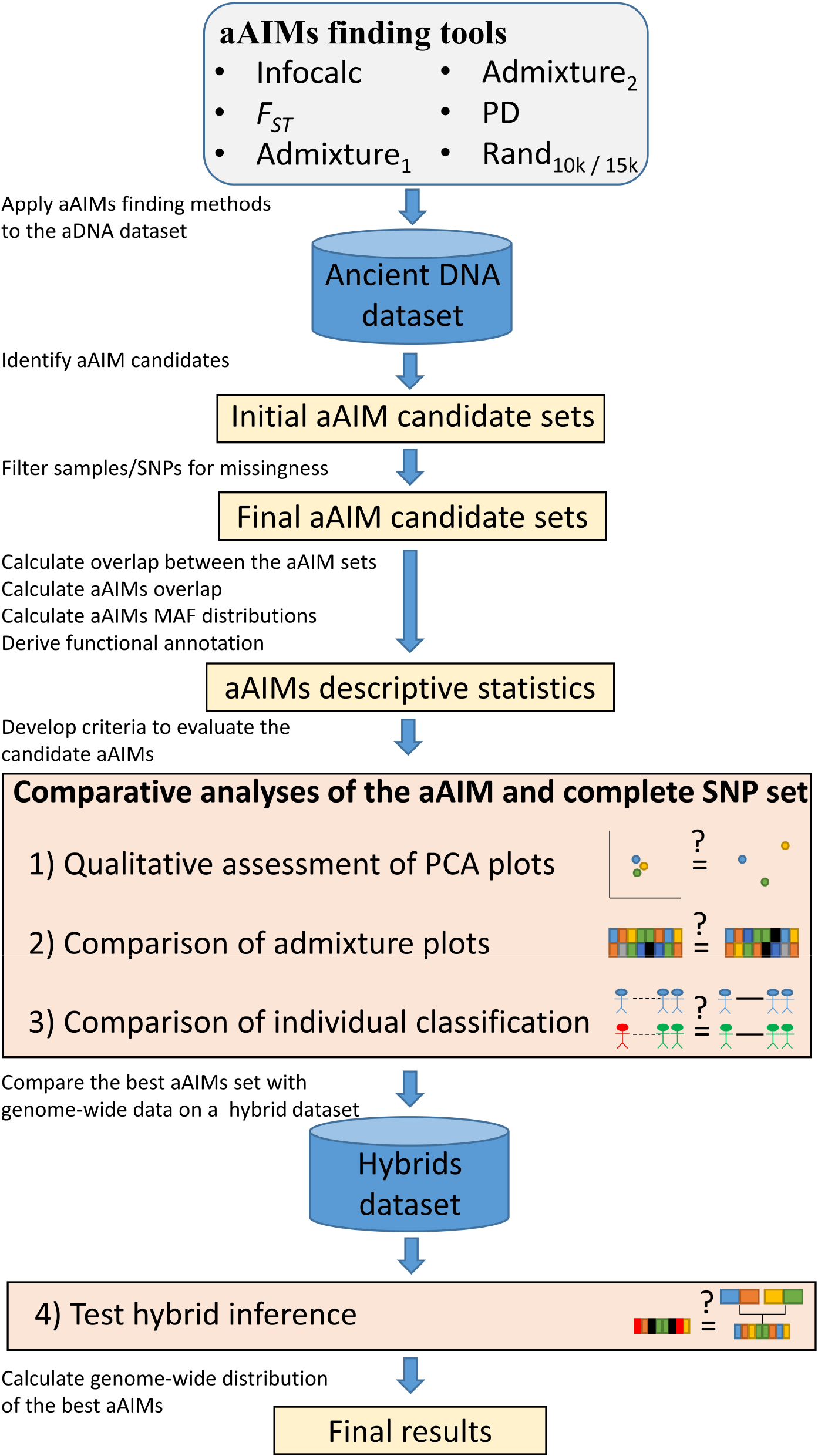
Our scheme to identify and evaluate the accuracy of aAIM finding algorithms compared to each other and to the complete SNP set. We adopted four criteria to evaluate how well the aAIMs candidate capture the population structure depicted by the complete SNP set (CSS): first, by qualitatively comparing the dispersal of genomes obtained from a PCA to that of the CSS. Second, by comparing the Euclidean distances between the admixture proportions of each genome and those obtained from the CSS. To avoid inconsistencies between the SNP sets, we used admixture components obtained through a *supervised* ADMIXTURE (see *methods*). Third, by testing which aAIMs classify individuals to populations most accurately. Finally, the ability to identify admixed individuals was evaluated for the top performing method against the CSS.

## Results

### Depicting ancient population structure

We constructed a dataset of 150,278 autosomal SNPs from 302 ancient genomes classified into 21 populations from Europe, the Middle East, and North Eurasia and dated to time periods spanning 14,000 years ago through 1,500 years ago (Figure 3, Table S1). Due to the limited availability of ancient genomes, our dataset was not uniform over time and space. For instance, there were 57 Central European genomes from the Late Neolithic to the Bronze Age, but populations such as Mesolithic Central and Western Europeans, Bronze Age Jordanians, Chalcolithic Russians, and Mesolithic Russians, comprised of three genomes each.

**Figure 3.**
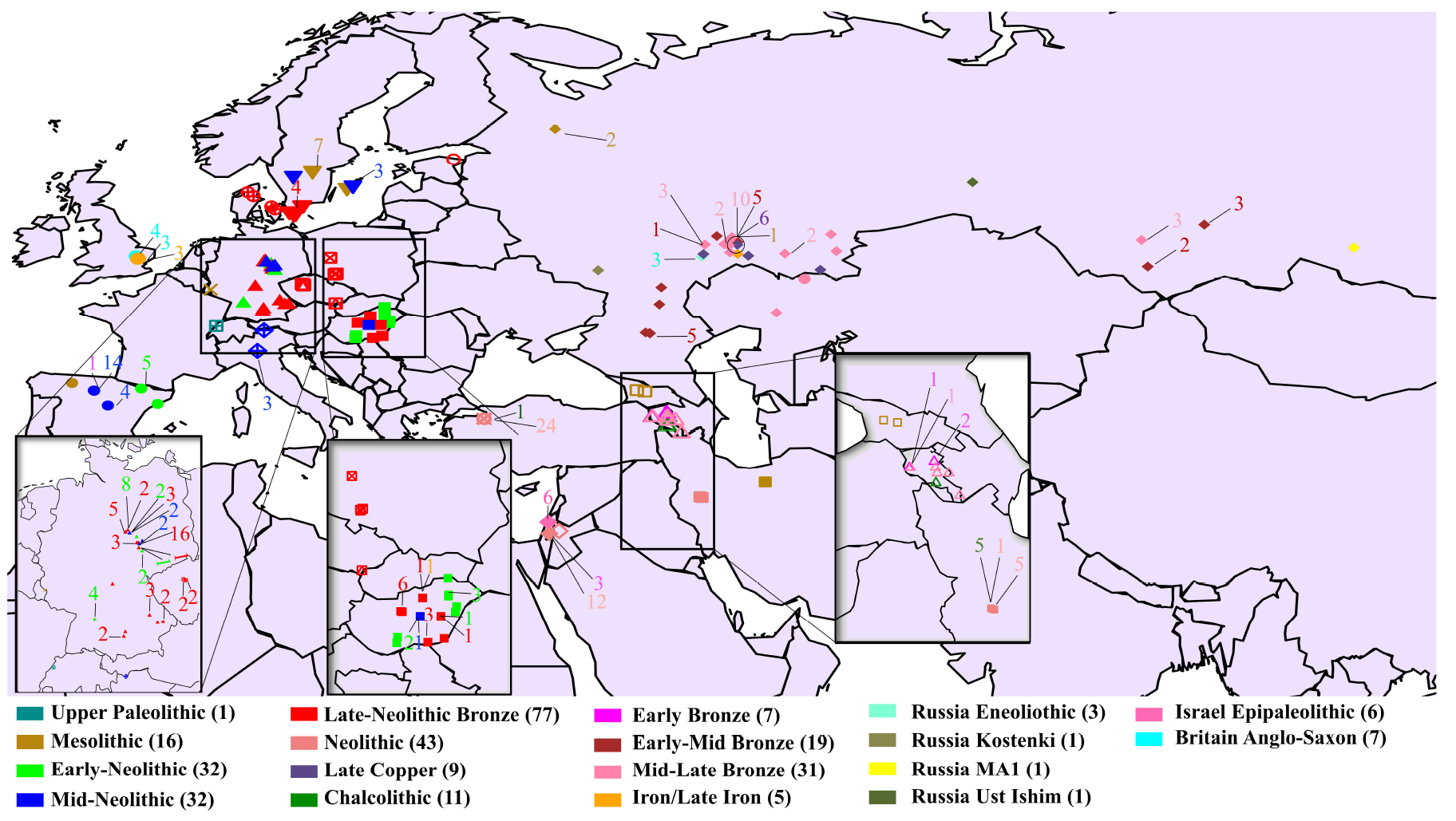
Geographical locations of the ancient genomes. The geographical coordinates of the ancient genomes. The shapes plotted in the map designate the country of origin of the genomes and their colors designate the era. The total number of ancient genomes from a specific era is shown.

Missingness varied greatly within the samples, as well as within the markers. The sample-based missingness ranged from 0.05% (KK1) to 99.2% (I1951) with an average of 54%. Similarly, missingness also varied among the populations, with Levantine Epipaleolithic and Neolithic genomes having the highest missingness (*n*=19, *μ*=90±16%) and Mesolithic Swedish genomes having the lowest (*n*=8, *μ*=29±27%). The SNP-based missingness ranged from 30% to 98% with an average of 54%.

Principal component analysis (PCA) of the ancient genomes substantiated previous observations of a Europe-Middle East contrast along the vertical principal component (PC1) and parallel clines (PC2) in both Europe and the Middle East (Figure S1). Genomes from the Epipaleolithic and Neolithic Levantine clustered at one extreme of the Near East-Europe cline with some overlapping with Neolithic Turkish and Central European genomes. Neolithic Iranians clustered between Central European genomes. While ancient Spanish, Armenian, Central EU, and British genomes occupied the intermediate position of Near Eastern and North Eurasian genomes, Russian and Swedish genomes clustered at the end of the Near East-Europe cline.

Our *unsupervised* ADMIXTURE analysis with various number of splits (*K*) (Figure S2) found that no choice of *K* minimized the cross-validation error (CVE) (Figure S3), likely because the high noise and missingness in the data prevented the CVE from stabilizing. At *K*=10 (Figure S4) multiple genomes (e.g., Britain Iron Saxon, Mesolithic Neolithic Caucasus population, Bronze Age Jordanian and Epipaleolithic Levantine, Chalcolithic, Mesolithic and Early Mid Bronze Russian, Early Neolithic Spanish, Mesolithic and Mid Neolithic Swedish, and Neolithic Turkish) appeared to be homogeneous in relation to their population and exhibited a distinct allelic frequency profile of admixture components. For these reasons, we decided to choose *K*=10 as the optimal value. Furthermore, in this case, putative ancient ancestral components, such as the *Early Neolithic European* (brown) and the *Russia Mid Late Bronze* (magenta), predominantly found among European genomes, may be identified. Except for their predominance in Neolithic Turkish genomes, these two components also exist in most Neolithic Central Europeans. Some 20-30% of Central European genomes have discernible fractions of *Europe Late Neolithic-Early Bronze* (navy-blue) and *Russia Mid-Late Bronze* (deep-pink) components, respectively. Two components (cyan and dark purple) appeared sporadically in a few populations, likely due to noise.

### Identifying and describing the aAIM candidates

We developed a framework to identify and evaluate the efficacy of aAIM candidates in capturing ancient population structure and allowing admixture mapping (Figure 2). aAIM candidates were identified using five methods (Figure 2). Similarly to the CSS, genomes and SNPs with over 90% missingness were removed, leaving each dataset with 223-263 genomes (Table S2). Furthermore, 310 SNPs without data were removed from the Rand_10k_ dataset. The final number of aAIM candidates is shown in Table S3. Overlapping aAIMs between the methods are remarkably small and range from 560 (Rand_10k_ and Admixture_1_ to 2,160 (Admixture_1_ and Admixture_2_). Interestingly, Infocalc and *F*_*ST*_, oftentimes used together, share only ~10% of their aAIM candidates. The PD method shares 13.7% of its aAIMs with *F*_*ST*_ and ~10% with Infocalc.

Comparing the properties of the aAIM candidates (Figure S5a), we found that Infocalc prioritized SNPs with the lowest MAF (45% of the aAIMs have MAF<0.1) and *F*_*ST*_ captured aAIM with high frequency of low-mid MAF. By contrast, PD and the Admixture-based methods exhibited higher frequencies of high MAF SNPs, with Admixture_2_ having the highest proportion of high MAF aAIMs (75% of the aAIMS have MAF>0.4). Interestingly, the MAF distributions exhibited similarity with modern populations (Figure S5b), though, for all methods, with fewer alleles in the lowest MAF bins. Unsurprisingly, most of the aAIM variants were non-functional (94.6-96.3%) and vary little from the CSS’s annotation (Table S4).

### Comparative testing of aAIM candidates

The accuracy of the aAIMs was evaluated using four criteria and comparing each method against both CSS and two random SNP sets of sizes that approximated the number of aAIM candidates. We first calculated the PCA for each SNP set and compared the population dispersion along the primary two axes. Similarly to the CSS (Figure S1), all the methods depicted the Europe-Middle East contrast (PC1) and parallel clines (PC2) in the European genomes so that ancient Jordanian, Levantine, Turkic, and Spanish genomes clustered at one extreme of the Near East-Europe cline, whereas the genomes from Russia and Sweden clustered at the other end (Figure S6). However, much like the random sets, Infocalc and *F*_*ST*_ did not separate Levantine and Turkic individuals from each other. Infocalc also merged the Caucasus individuals with central Europeans. The admixture-based methods and PD separated all the ancient populations, similarly to the CSS and better, in the case of Scandinavians and Russians.

We next quantitatively assessed which dataset produced the closest admixture signature to that of the CSS (Figure S4). For that, we calculated the admixture proportions in relation to the ten putatively ancient ancestral populations that we obtained with the CSS (Figures S7-8) and then computed their Euclidean distances to their counterparts obtained with the CSS (Figure 4). The PD aAIMs led to significantly shorter Euclidean distances (*μ*=0.13, *σ*= 0.1, *n*=302) compared to those obtained from the other aAIMs (Welch *t*-test *p*-values: Infocalc 0.002, *F*_*ST*_ 8.5×10^−13^, Admixture_1_ 2.2×10^−16^, Admixture_1_ 2×10^−16^, Rand_10k_ 5×10^−6^, and Rand_15k_ 0.001). Infocalc’s aAIMs produced the second shortest distances from the CSS (*μ*=0.17, *σ*=0.15); however, they were not statistically shorter than the distances obtained with the two random datasets (Welch *t*-test *p*-values: Rand_10k_ 0.12 and Rand_15k_ 0.77 respectively), suggesting that Infocalc was unable to capture the population structure. *F*_*ST*_-derived AIMs (*μ*=0.2, *σ*=0.13) performed worse than the Rand_15k_ aAIMs (Welch *t*-test *p*-value 0.004), and similarly to the Rand_10k_ aAIMs (Welch *t*-test, *p*-value=0.13). Finally, the two admixture-based datasets performed the worst performances of all the methods (*μ*_1_=0.22, *σ*_*1*_=0.15 and *μ*_*2*_=0.24, *σ*_*1*_=0.16), being even significantly worse than the two random datasets (Welch t-test: Admixture_1_ [Rand_10k_ *p*-value=0.002] and [Rand_15k_ *p*-value=1.6×10^−5^]; Admixture_2_ [Rand_10k_ *p*-value=1.7×10^−5^] and [Rand_15k_ *p*-value=2.5×10^−8^]).

**Figure 4.**
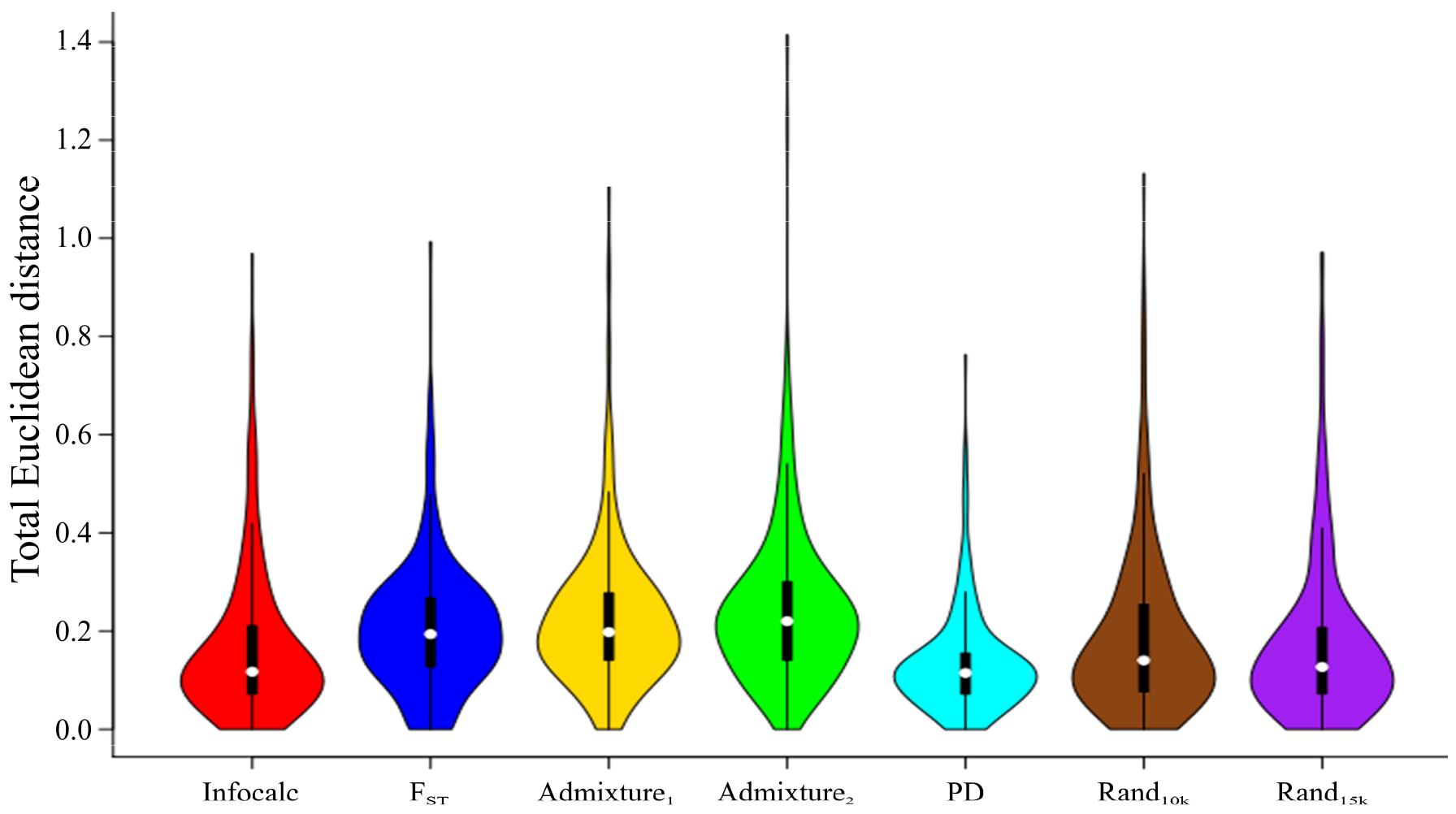
A comparison of the Euclidean distances (Δ) between the admixture proportions of the ancient genomes obtained from the CSS and those obtained from the aAIM sets. Lower distances indicate high genetic similarity between the admixture proportions obtained using two different SNP sets.

Third, we assessed which aAIMs dataset allows classifying individuals into population groups most accurately. An admixture-based population classifier was applied to the admixture proportions produced by all the datasets, and their accuracy was compared to that of the CSS (76±25%) and the known population classification (Table S1). The mean classification accuracy per population ranged from 3% (*F*_*ST*_) to 61% (PD) with the PD outperforming all the other methods (Table 1). In other words, ~13k (8%) of the SNPs are sufficiently informative to classify individuals to populations with 80% of the accuracy of the CSS. In nine out of 21 population groups (22% of the individuals) PD-based classification was similar or more accurate than the CSS. All other methods performed similarly or worse than the random SNP sets (42±22% and 50±23%) with Infocalc (50±23%) outperforming the remaining methods. Of note is the poor performance of *F*_*ST*_ aAIMs, likely due to the high sensitivity of *F*_*ST*_ to aDNA data. As expected, high missingness was associated with incorrect predictions (Figure S9). For example, the low-coverage, low-quality Britain Anglo-Saxon genomes proved challenging for all the methods (0-40%) but predicted correctly with the CSS (100%).

**Table 1.**
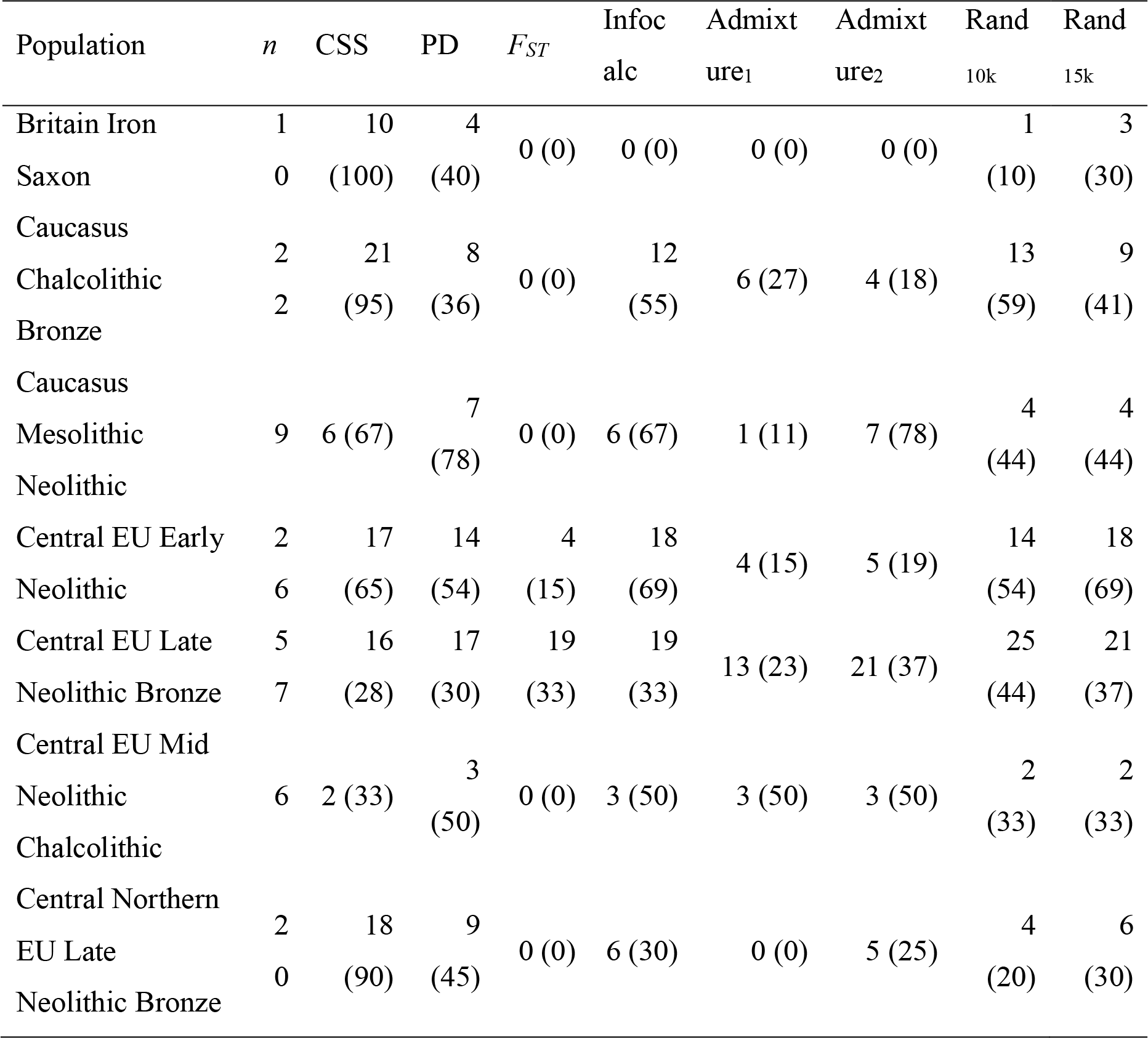
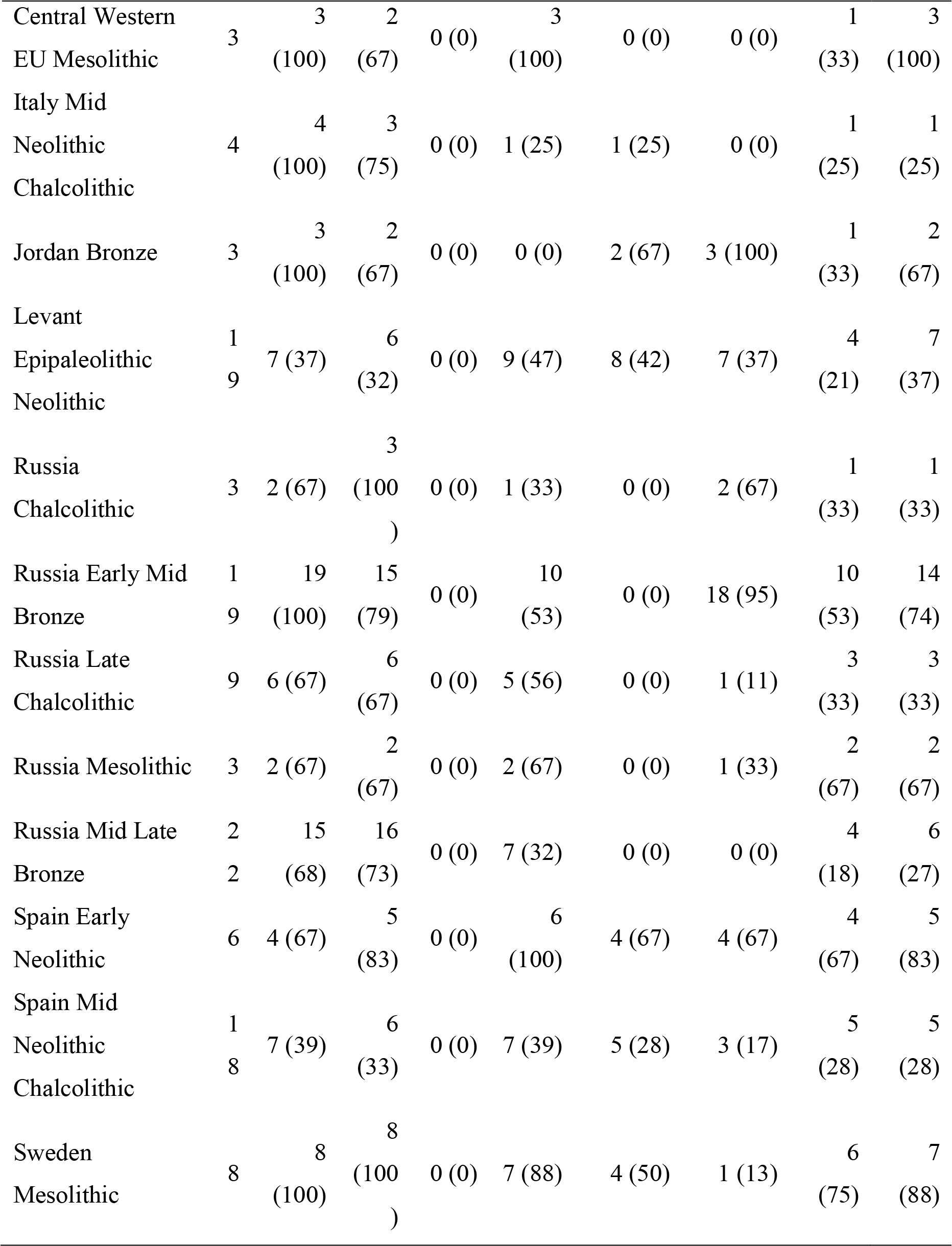
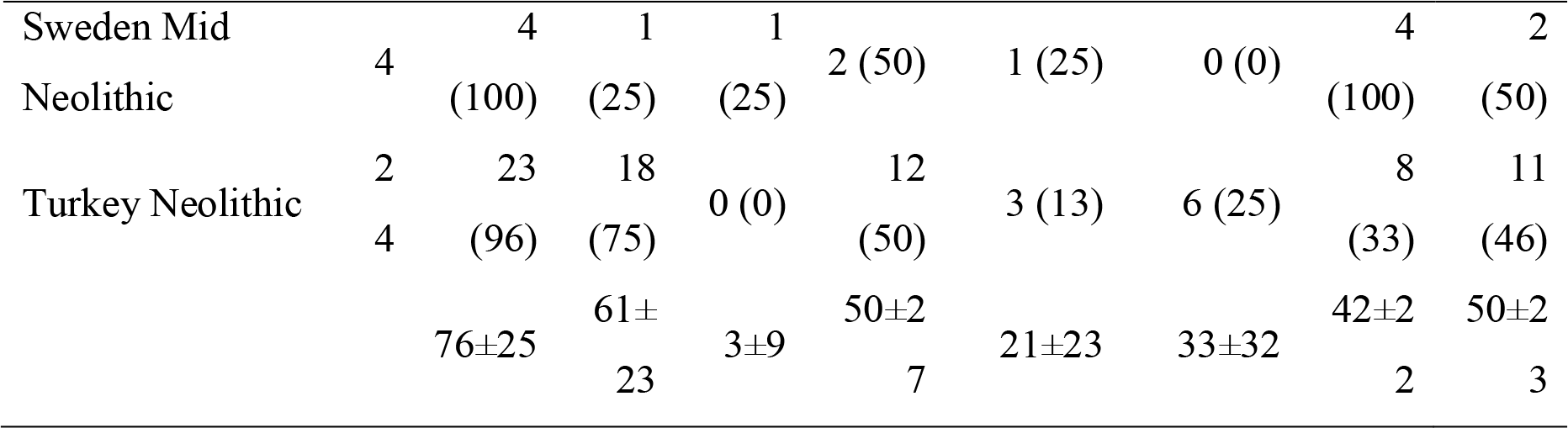
Accuracy in classifying individuals to populations using the aAIM candidates. Each column reports the number of individuals correctly predicted and, in brackets, the corresponding population percentage. The total number of individuals per population are reported in the second column. Mean and standard deviation for each SNP set are provided in the last row.

### Inference of admixed samples

The last criterion used to evaluate the accuracy of the aAIMs was to test whether they can identify hybrid individuals. Due to the high accuracy of the PD aAIMs in classifying individuals into populations, when compared to the alternative datasets, we then decided to drop the other methods and tested only the aAIMs derived from PD. Figure S10 illustrates the genome-wide distribution of PD aAIMs. To assess whether these aAIMs can identify hybrid individuals, ancient individuals were hybridized to form 120 mixed individuals, each associated with three datasets: CSS, PD aAIMs, and a random SNP set of the size of PD aAIMs (Table 2).

**Table 2.** Accuracy of inferring hybrid individuals using the PD aAIMs. The parental populations and the number of hybrids generated from them are shown. Each hybrid was represented by three datasets: CSS, PD aAIMs, and a random SNP set. The average genetic distances (d) between the admixture components of these datasets per population are shown.

| Parental<br>population<br>A | Parental<br>population<br>B | #<br>Hybrids | $\overline{d}(\text{CSS}, \text{PD})$ | $\overline{d}(\text{CSS}, \text{random set})$ | $\overline{d}(\text{PD}, \text{random set})$ |
| --- | --- | --- | --- | --- | --- |
| Britain Iron<br>Saxon | Britain Iron<br>Saxon | 6 | 0.026 | 0.212 | 0.208 |
| Britain Iron<br>Saxon | Russia Late<br>Chalcolithic | 9 | 0.009 | 0.610 | 0.601 |
| Britain Iron<br>Saxon | Sweden<br>Mesolithic | 9 | 0.051 | 0.344 | 0.337 |
| Britain Iron<br>Saxon | Turkey<br>Neolithic | 9 | 0.003 | 0.428 | 0.431 |
| Britain Iron<br>Saxon | Spain Early<br>Neolithic | 9 | 0.108 | 0.221 | 0.241 |
| Russia Late<br>Chalcolithic | Russia Late<br>Chalcolithic | 6 | 0.009 | 0.443 | 0.448 |
| Russia Late<br>Chalcolithic | Sweden<br>Mesolithic | 9 | 0.062 | 0.578 | 0.561 |
| Russia Late<br>Chalcolithic | Turkey<br>Neolithic | 9 | 0.063 | 0.661 | 0.633 |
| Russia Late<br>Chalcolithic | Spain Early<br>Neolithic | 9 | 0.101 | 0.520 | 0.491 |
| Sweden<br>Mesolithic | Sweden<br>Mesolithic | 6 | 0.000 | 0.384 | 0.384 |
| Sweden<br>Mesolithic | Turkey<br>Neolithic | 9 | 0.055 | 0.567 | 0.522 |
| Spain Early<br>Neolithic | Sweden<br>Mesolithic | 9 | 0.108 | 0.402 | 0.377 |
| Turkey<br>Neolithic | Turkey<br>Neolithic | 6 | 0.001 | 0.627 | 0.626 |
| Spain Early<br>Neolithic | Turkey<br>Neolithic | 9 | 0.092 | 0.483 | 0.493 |
| Spain Early<br>Neolithic | Spain Early<br>Neolithic | 6 | 0.041 | 0.197 | 0.172 |

The genetic admixture distances between the admixture components generated using the CSS and PD aAIMs were significantly smaller (*μ*=0.05, *σ*=0.04) than both the genetic admixture distances between the CSS and the random SNP set (*μ*=0.45, *σ*=0.15, Welch *t*-test *p*-values=2.2×10^−8^) and those between the PD and the random SNP set (*μ*=0.43, *σ*=0.15, Welch *t*-test *p*-values=1.9×10^−8^). We thus demonstrated that PD aAIMs can be used to study admixed individuals and for future admixture mapping involving aDNA.

## Discussion

Questions of identity are at the center of scientific and public debate. Until recently, charting the emergence of agriculture, the spread of languages, and the rise and decline of cultures were topics dominated by archeologists. The emergence of aDNA allows paleogeneticists to delve into this debate with a discordant set of assumptions about biology and identity [30]. This was not unforeseen, as population genetic analyses excel at identifying individual differences, which can inform archeologically contended subjects like migration and the degree of admixture or population replacements. However, aDNA analyses also require destroying genetic material, sometimes irrevocably, which makes them impossible to replicable. It is thereby crucial to develop a robust genetic methodology that uses population genetic principles to examine the assumptions made by both archeologists and paleogeneticists. It is reasonable to expect that many of the tools employed to study modern-day genomes will need to be adapted to the four-dimensional environment facilitated by aDNA.

Ancestry informative markers (AIMs) are some of the most useful tools in addressing population, biomedical, forensics, and evolutionary questions that remain in use today [8, 31–33]. However it is unclear to what extent known AIMs are applicable to ancient genomic data, characterized by high missingness and haploidy[1].

In this study, we defined ancient ancestry informative markers (aAIMs) (Figure 1) and sought to identify them using various methods. The number of aAIM candidates detected by each method ranges from 9,000 to 15,000. These numbers are of the same magnitude as large AIMs studies [e.g., 34, 35] and reasonable, provided the potential relatedness of the ancient Eurasian populations and the near absence of heterozygote markers in the data. To find which of the aAIM candidate sets produced by each method best represent the true population structure, we used the CSS as a benchmark for qualitatively and quantitatively comparisons.

Identifying the ideal AIM set that would be both small and include redundancies (in case of sequencing failure), capture the population structure, and allow identifying admixed individuals is one of the challenges of population genetics. We showed that aAIMs identified through a PCA-derived (PD) method outperformed all other methods, in agreement with previous studies that tested PCA-based methods [24]. Forty percent of the classifications made by the PD method were more accurate than those made using the CSS, which highlights the limitations of using markers indiscriminately. This is not surprising, since not all markers are equally informative and less-informative markers (e.g., exomic markers) may mask the population structure resulting in misclassification of populations. The notion of “more is better” is thereby particularly misguided with aDNA that harbors multi-layered population structure in a poor set of markers. The applicability of the PD aAIMs for admixture mapping combined with tools that can homogenize cases and controls [e.g., 15] allows for carrying out future association studies on aDNA samples [e.g., 3]. Further investigations with additional data may identify formerly common markers associated with the disease that with time became rare and undetectable.

Surprisingly, Infocalc and *F*_*ST*_ aAIMs, typically used in conjunction to identify AIMs [17] and reported to perform well [36], have oftentimes underperformed random SNP selections. Not only was *F*_*ST*_ already shown to be particularly small within continental populations [37], but these methods may be particularly sensitive to aDNA data that is both haploid and has high missingness (Figure S9). We also found no relationships between MAF and aAIMs performances (Figure S5). Enrichment for high or low MAF SNPs did not guarantee success, although the PD harbored more common SNPs than most underperforming methods.

Our study has several limitations. We studied an uneven number of Eurasian populations from various times and locations, causing a skew towards markers that predict central European populations from the Late Neolithic and Bronze Age. A modest attempt to reduce this bias was made by including modern-day African and Asian populations; however, a more comprehensive analyses should be made when more global genomes are available from consecutives eras. Second, the aAIMs were calculated independently by each method with individual populations considered independent, although the PCA and ADMIXTURE plots indicate that central European populations may not be independent. Finally, due to high missingness of the data, it is likely that our study missed informative markers that could improve the classification accuracy in newly sequenced populations. Our framework and methods should therefore be applied again when more comprehensive aDNA datasets are available.

## Conclusions

The use of ancient genomes in research is in its infancy and expected to intensify and expand to new fields as more data become available. One of the main advantages of aDNA is that it widens the number of ancestry types and makes them multi-faceted, requiring fine-tuned molecular utilities to depict ancestry over time. AIMs are some of the most effective tools that have spear-headed population genetics over the past two decades and are ancillary to the challenge of understanding population structure. Here, we defined ancient AIMs (aAIMs), proposed a framework to evaluate AIM-finding methods, demonstrated the accuracy of a novel aAIM-finding method, and reported the most successful set of aAIMs. Future analyses may benefit from using our framework, methods, and aAIMs in order to refine ancient population structure models.

## Methods

### Data collection

Genomic data were obtained from 11 publications depicting 302 ancient genomes (Table S1). In the case of sequence data, sequence reads were aligned to the human reference assembly (UCSC hg19-http://genome.ucsc.edu/) using the Burrows Wheeler Aligner (BWA version 0. 7.15) [38], allowing two mismatches in the 30-base seed. Alignments were then imported to binary (bam) format, sorted, and indexed using SAMtools (version 1.3.1) [39]. Picard (version 2.1.1) (http://picard.sourceforge.net/) was then used for MarkDuplicates to remove reads with identical outer mapping coordinates (which are likely PCA artifacts). The Genome Analysis Toolkit RealignerTargetCreator module (GATK version 3.6) [40, 41] was used to generate SNP and small InDel calls for the data within the targeted regions only. GATK InDelRealigner/BaseRecalibrator was then used for local read realignment around known InDels and for base quality score recalibration of predicted variant sites based on dbSNP 138 and 1000 Genomes known sites, resulting in corrections for base reported quality. The recalibration was followed by SNP/InDel calling with the GATK HaplotypeCaller. Variants were filtered for a minimum confidence score of 30 and minimum mapping quality of 40. At the genotype level, all genotypes that had a genotype depth less than 4 (GD < 4) or a genotype quality score less than 10 (GQ < 10) were removed from the dataset by setting them to missing in the VCF. GATK DepthofCoverage was then used to re-examine coverage following the realignment. VCFtools (version 0.1.14) [42] were used to convert the VCF file to PLINK format [43]. The final dataset comprised of 150,278 autosomal SNPs from 302 ancient DNA (aDNA) genomes (Table S1; Additional file 1). Eight aDNA genomes (I0247, I1584, ATP9, IR1, Kostenki14, MA1, and Ust Ishim) without any country/region designation were omitted in the closest population determination calculations. The genomes were divided into 21 populations, based on their sampling country/region and era.

### Data analyses

#### The genetic structure canvas of ancient Eurasian genomes

The population structure of the ancient genomes was described using principal component analysis (PCA) implemented in PLINK v1.9 [43]. Genomes and SNPs with over 90% missingness were removed. We also applied the model-based clustering methods implemented in ADMIXTURE v1.3 [44]. Minor allele frequency (MAF) was calculated using PLINK (-maf command). For modern-day populations, MAF was calculated from the 1000 Genomes populations (ALL.2of4intersection.20100804.genotypes) [45]. Percentage of rare and novel variants and other functional information were obtained through VEP.

#### Identifying aAIMs

We applied two established and three novel methods to detect aAIM candidates:

1. Infocalc v1. 1 [27], which determines the amount of information multiallelic markers provide about an individual’s ancestry by calculating the informativeness (*I*) of each marker separately and ranking the SNPs by their informativeness. Infocalc determines *I* based on the mathematical expression described in Rosenberg et al. (2003). We compared the performances (Figure 2) of the top 5,000, 10,000, 15,000, and 20,000 most informative markers (results not shown). The 15,000 dataset outperformed all other datasets and was selected for further analyses.
2. *F*_*ST*_. Wright’s fixation indices (*F*_*ST*_) [29] measures the degree of differentiation among populations potentially arising due to genetic structure within populations. Given a set of populations (Table S1), we employed PLINK [43] to estimate *F*_*ST*_ separately for all the markers using ‒fst command alongside --within flag. Due to the high fragmentation of the data, *F*_*ST*_ values could only be calculated for 46% of the dataset. We compared the performances (Figure 2) of 5,000, 10,000, 15,000, and 20,000 SNPs with the highest *F*_*st*_ values (results not shown). The 15,000 dataset outperformed all other datasets and was selected for further analyses.
3. Admixture_1_. This method assumes that aAIMs have high allelic frequencies in certain subpopulations that drive the differentiation of admixture components. Analyzing ADMIXTURE’s output file (P file) for *K*=10, we identified the markers (rows) that had high allele frequency (>0.9) in only one admixture component (columns). Comparing the number of high-MAF SNPs in all columns, we selected 9,309 from the five columns with the highest number of such SNPs.
4. Admixture_2_. This method assumes that aAIMs embody both high allelic frequencies in certain subpopulations and high variance between these allelic frequencies that differentiate admixture components. Analyzing ADMIXTURE’s output file for *K* of 10, we identified 11,418 SNPs showing high variance (≥0.04) and high allele frequency range (maxima - minima ≥ 0.65) between the admixture components.
5. PC-based (PD) approach. This method assumes that AIMs can replicate the population structure of subpopulations represented by the variation in the first two PCs. This is an interactive PC-based approach that identifies the smallest set of markers necessary to capture the population structure of a group of individuals as captured by the CSS. More specifically, for each population group (Table S1) in which at least 100 SNPs were available, we calculated PCA and used PC1 and PC2 to plot the individuals after all SNPs with high missingness (>0.05) were excised. If the population group had insufficient SNPs, we relaxed the missingness threshold by an additional 0.05, though 0.05 were sufficient for almost all groups. We then scored the SNPs by their informativeness, as in [46], and visually compared the plot to that obtained from the CSS (Figure S11). If the plots were dissimilar, we repeated the analysis using additional 100 top scored SNPs until either the plots exhibited high similarity, or a threshold of 2000 SNPs was reached. We were unable to complete the analyses for 3 populations due to the small number of individuals. The PD method is available on https://github.com/eelhaik/PCA-derived-aAIMs. On average, 861 SNPs were collated per population group. Overall, the dataset comprised of 13,027 SNPs.

To compare the prediction accuracy of the aAIMs subsets, two control datasets (Rand_10k_ and Rand_15k_) were generated by randomly sampling 10,000 and 15,000 SNPs from the CSS, respectively. aAIMs identified by all methods are available as Additional file 2.

#### Classifying individuals into populations from genomic data

***Identifying ancient admixture components***. We selected 100 random ancient genomes and removed six for insufficient data (>95% missingness). To those, we added 20 Han Chinese and 20 Yoruba modern genomes from the 1000 Genomes Project (Durbin et al. 2010). We then applied *supervised* ADMIXTURE with various *K*’s ranging from 8 to 13. While we were unable to find a single *K* where culturally related genomes exhibited homogeneous admixture patterns, the most robust population substructure was found for *K* of 10. Two more components were obtained by analyzing Spanish and German genomes that appeared indistinguishable along with five Yoruba genomes separately. We observed very little admixture with the Han and Yoruba. Overall, we identified 10 admixture components in ancient genomes, corresponding to the allele frequencies of 10 hypothetical populations. Similarly to Elhaik et al. [16], we simulated 15 samples for each hypothetical population by generating 30 alleles which MAF corresponds to the MAF of each population and assigning those genotypes to the simulated individuals. The putative ancestral ancient populations are available in Additional file 3. ***Relabeling populations***. Initially, the labels from the corresponding papers were used to classify individuals to population. The consistency of these labels with data was evaluated by carrying out a *supervised* ADMIXTURE analysis on the genomic data combined with the 150 putative ancient ancestral individuals. Due to the high similarity of the admixture patterns between individuals of different groups living in similar periods or entire groups (e.g., Neolithic individuals from Hungary and those from Germany), we re-labeled some of the population to reduce the number of populations and create more genomically homogeneous populations. For instance, Natufian and Neolithic samples from Jordan are grouped into the label Levant Epipaleolithic Neolithic. Overall, we identified 21 populations, whose labels are of the form “area_time period.” In the case of the Caucasus label, all the samples from Iran (except Iran_HotuIIIb) were excavated in the Zagros Mountains, south of the Caucasus. Given their admixture similarity with Armenians and Georgians from the same periods and their proximity to the Caucasus, this area was labelled as Caucasus. Iran_HotuIIIb was found in a more eastern region, just below the southeastern edge of the Caspian Sea, and given its similarity to Georgians and other Iranians, it was included in the group Caucasus Mesolithic Neolithic. ***Genomically defining reference populations*.** For each population with *N*_*P*_ > 4, where *N*_*P*_ is the number of individuals in the population, individuals were grouped in clusters through an iterative process that uses a *k*-means clustering technique paired with multiple pairwise *F*-tests. Iterations ran over the number of *k* clusters [2, *N*_*P*_/2]. At each iteration *i*, *k*-means was used to identify the *k* clusters, then the *F*-test was applied on each pair of clusters to test whether they are significantly (*P*<0.05) different. If the two clusters are different from all the pairs at iteration *i*, the process advances to *i*+1 until at least one pair violates the condition, in which case *k*_*op*_=*i*−1 is the optimal number of clusters or reference populations within that population. ***Assigning individuals to populations*.** We developed an admixture-based classifier, which is not sensitive to exclusion of random groups of individuals nor inclusion of large numbers of individuals from admixed groups and was trained on a third of the data. Using *supervised* ADMIXTURE, we calculated the admixture proportions of the individuals in relation to the putative ancient ancestral populations. Population assignment was then made based on the minimal Euclidean distance between the admixture components of each genome and those of the reference populations. The assignment accuracy was measured against the population classification (Table S1).

#### Assessing admixture mapping

***Creating hybrid individuals***. We selected 15 individuals from five populations that showed homogeneity in their admixture components (Figure S4) and randomly sampled 120 pairs. Since selecting random alleles from each parent was problematic due to the high missingness of the data, we randomly selected half the genotypes of each parent to form 120 “offspring” or hybrid genomes. Each hybrid had three SNP sets: the CSS, PD aAIMs, and a random SNP set of the size of PD aAIMs with SNPs selected randomly for each hybrid. ***Assessing admixture accuracy***. Following [47–49], we applied a *supervised* ADMIXTURE to the three SNP sets of each hybrid and calculated their genetic admixture distances (*d*) from each another, defined as the Euclidean distance between two set of admixture proportions.

#### Graphics

All PCA and Admixture plots were generated in R v3.2.3. Maps were drawn using the ‘rworldmap’ package implemented in R v3.2.3.

#### Availability of data and materials

The dataset supporting the conclusions of this article is included within the article and its additional files.

## Acknowledgment

This study was partially supported by the MRC Confidence in Concept Scheme award 2014-University of Sheffield to E.E. (Ref: MC_PC_14115). We thank Grace Holland who was partially supported by the UK EPSRC Doctoral Training Partnership Grant EP/N509735/1 as a Vacation Bursary Training Project.

## Supporting Information Legends

Elhaik et al 2018 – Supp – Figures S1-S11 and Tables S1-S4

Additional file 1.zip – Genotype data of the aDNA samples

Additional file 2.zip – aAIMs candidates used in all analyses

Additional file 3.zip – Genotype file of the putative ancient ancestral populations

## References

1. Morozova I, Flegontov P, Mikheyev AS, Bruskin S, Asgharian H, Ponomarenko P, Klyuchnikov V, ArunKumar G, Prokhortchouk E, Gankin Y, et al: Toward high-resolution population genomics using archaeological samples. DNA Res 2016, 23:295–310.

2. Marciniak S, Perry GH: Harnessing ancient genomes to study the history of human adaptation. Nat Rev Genet 2017, advance online publication.

3. Cassidy LM, Martiniano R, Murphy EM, Teasdale MD, Mallory J, Hartwell B, Bradley DG: Neolithic and Bronze Age migration to Ireland and establishment of the insular Atlantic genome. Proc Natl Acad Sci U S A 2016, 113:368–373.

4. Patterson NJ, Moorjani P, Luo Y, Mallick S, Rohland N, Zhan Y, Genschoreck T, Webster T, Reich D: Ancient admixture in Human history. Genetics 2012, 192:1065–1093.

5. Mathieson I, Lazaridis I, Rohland N, Mallick S, Patterson N, Roodenberg SA, Harney E, Stewardson K, Fernandes D, Novak M, et al: Genome-wide patterns of selection in 230 ancient Eurasians. Nature 2015, 528:499–503.

6. Fu Q, Hajdinjak M, Moldovan OT, Constantin S, Mallick S, Skoglund P, Patterson N, Rohland N, Lazaridis I, Nickel B, et al: An early modern human from Romania with a recent Neanderthal ancestor. Nature 2015.

7. Li JZ, Absher DM, Tang H, Southwick AM, Casto AM, Ramachandran S, Cann HM, Barsh GS, Feldman M, Cavalli-Sforza LL, Myers RM: Worldwide human relationships inferred from genome-wide patterns of variation. Science 2008, 319:1100–1104.

8. Elhaik E, Yusuf L, Anderson AIJ, Pirooznia M, Arnellos D, Vilshansky G, Ercal G, Lu Y, Webster T, Baird ML, Esposito U: The Diversity of REcent and Ancient huMan (DREAM): a new microarray for genetic anthropology and genealogy, forensics, and personalized medicine. Genome Biol Evol 2017, 9:3225–3237.

9. Elhaik E, Greenspan E, Staats S, Krahn T, Tyler-Smith C, Xue Y, Tofanelli S, Francalacci P, Cucca F, Pagani L, et al: The GenoChip: a new tool for genetic anthropology. Genome Biol Evol 2013, 5:1021–1031.

10. Hublin J-J: The last Neanderthal. Proc Natl Acad Sci USA 2017:201714533.

11. Jones S: The archaeology of ethnicity: constructing identities in the past and present. London: Routledge; 1997.

12. Albrechtsen A, Nielsen FC, Nielsen R: Ascertainment biases in SNP chips affect measures of population divergence. Mol Biol Evol 2010, 27:2534–2547.

13. Marchini J, Cardon LR, Phillips MS, Donnelly P: The effects of human population structure on large genetic association studies. Nat Genet 2004, 36:512–517.

14. Yusuf S, Wittes J: Interpreting geographic variations in results of randomized, controlled trials. N Engl J Med 2016, 375:2263–2271.

15. Elhaik E, Ryan D: Pair Matcher (PaM): fast model-based optimisation of treatment/case-control matches using demographic and genetic data. bioRxiv 2017.

16. Elhaik E, Tatarinova T, Chebotarev D, Piras IS, Maria Calò C, De Montis A, Atzori M, Marini M, Tofanelli S, Francalacci P, et al: Geographic population structure analysis of worldwide human populations infers their biogeographical origins. Nat Commun 2014, 5:1–12.

17. Phillips C, Parson W, Lundsberg B, Santos C, Freire-Aradas A, Torres M, Eduardoff M, Borsting C, Johansen P, Fondevila M, et al: Building a forensic ancestry panel from the ground up: The EUROFORGEN Global AIM-SNP set. Forensic Sci Int Genet 2014, 11:13–25.

18. Kosoy R, Nassir R, Tian C, White PA, Butler LM, Silva G, Kittles R, Alarcon-Riquelme ME, Gregersen PK, Belmont JW, et al: Ancestry informative marker sets for determining continental origin and admixture proportions in common populations in America. Hum Mutat 2009, 30:69–78.

19. Qin H, Zhu X: Power comparison of admixture mapping and direct association analysis in genome-wide association studies. Genet Epidemiol 2012, 36:235–243.

20. B. Bf, F. Cn, Milena S, A. Fa, R. Tf, C. Mm, K. At, L. Gdslv, A. De, L. Sa: Ancestry Informative Marker Panel to Estimate Population Stratification Using Genome-wide Human Array. Ann Hum Genet 2017, 81:225–233.

21. Qian P, J. Sn, C. Wk, L. Ec: Whole genome sequence association and ancestry-informed polygenic profile of EEG alpha in a Native American population. Am J Med Genet B Neuropsychiatr Genet 2017, 174:435–450.

22. Shriner D: Overview of admixture mapping. Curr Protoc Hum Genet 2013, Chapter 1:Unit 123.

23. Kidd KK, Speed WC, Pakstis AJ, Furtado MR, Fang R, Madbouly A, Maiers M, Middha M, Friedlaender FR, Kidd JR: Progress toward an efficient panel of SNPs for ancestry inference. Forensic Sci Int Genet 2014, 10:23–32.

24. Huckins LM, Boraska V, Franklin CS, Floyd JAB, Southam L, Gcan, Wtccc, Sullivan PF, Bulik CM, Collier DA, et al: Using ancestry-informative markers to identify fine structure across 15 populations of European origin. Eur J Hum Genet 2014, 22:1190.

25. Xu S, Huang W, Qian J, Jin L: Analysis of genomic admixture in Uyghur and its implication in mapping strategy. Am J Hum Genet 2008, 82:883–894.

26. Pakstis AJ, Kang L, Liu L, Zhang Z, Jin T, Grigorenko EL, Wendt FR, Budowle B, Hadi S, Al Qahtani MS, et al: Increasing the reference populations for the 55 AISNP panel: the need and benefits. Int J Legal Med 2017, 131:913–917.

27. Rosenberg NA, Li LM, Ward R, Pritchard JK: Informativeness of genetic markers for inference of ancestry. Am J Hum Genet 2003, 73:1402–1422.

28. Kidd JR, Friedlaender FR, Speed WC, Pakstis AJ, De La Vega FM, Kidd KK: Analyses of a set of 128 ancestry informative single-nucleotide polymorphisms in a global set of 119 population samples. Investig Genet 2011, 2:1.

29. Wright S: Evolution and the genetics of populations. A treatise in three volumes. Chicago, IL: University of Chicago Press; 1968.

30. Callaway E: Divided by DNA: The uneasy relationship between archaeology and ancient genomics. Nature 2018, 555:573–576.

31. Bose N, Carlberg K, Sensabaugh G, Erlich H, Calloway C: Target capture enrichment of nuclear SNP markers for massively parallel sequencing of degraded and mixed samples. Forensic Sci Int Genet 2018, 34:186–196.

32. Bulbul O, Speed WC, Gurkan C, Soundararajan U, Rajeevan H, Pakstis AJ, Kidd KK: Improving ancestry distinctions among Southwest Asian populations. Forensic Sci Int Genet 2018, 35:14–20.

33. Lopez-Cortes A, Echeverria-Garces G, Burgos G, Zambrano A, Cabrera-Andrade A, Garcia-Cardenas J, Salazar C, Leone P, Paz-y-Miño C: Molecular analysis of ancestry informative markers (AIMs-INDELs) in a high altitude Ecuadorian mestizo population affected with breast cancer. Forensic Science International: Genetics Supplement Series 2017, 6:e231–e232.

34. Tian C, Hinds DA, Shigeta R, Adler SG, Lee A, Pahl MV, Silva G, Belmont JW, Hanson RL, Knowler WC, et al: A genomewide single-nucleotide-polymorphism panel for Mexican American admixture mapping. Am J Hum Genet 2007, 80:1014–1023.

35. Paschou P, Lewis J, Javed A, Drineas P: Ancestry informative markers for fine-scale individual assignment to worldwide populations. J Med Genet 2010, 47:835–847.

36. Ding L, Wiener H, Abebe T, Altaye M, Go RC, Kercsmar C, Grabowski G, Martin LJ, Hershey GK, Chakorborty R, Baye TM: Comparison of measures of marker informativeness for ancestry and admixture mapping. BMC Genomics 2011, 12:622.

37. Elhaik E: Empirical distributions of Fst from large-scale Human polymorphism data. PLoS One 2012, 7:e49837.

38. Li H, Durbin R: Fast and accurate short read alignment with BurrowsWheeler transform. Bioinformatics 2009, 25:1754–1760.

39. Li H, Handsaker B, Wysoker A, Fennell T, Ruan J, Homer N, Marth G, Abecasis G, Durbin R: The Sequence Alignment/Map format and SAMtools. Bioinformatics 2009, 25:2078–2079.

40. McKenna A, Hanna M, Banks E, Sivachenko A, Cibulskis K, Kernytsky A, Garimella K, Altshuler D, Gabriel S, Daly M, DePristo MA: The genome analysis toolkit: a MapReduce framework for analyzing next-generation DNA sequencing data. Genome Res 2010, 20:1297–1303.

41. DePristo MA, Banks E, Poplin R, Garimella KV, Maguire JR, Hartl C, Philippakis AA, del Angel G, Rivas MA, Hanna M, et al: A framework for variation discovery and genotyping using next-generation DNA sequencing data. Nat Genet 2011, 43:491–498.

42. Danecek P, Auton A, Abecasis G, Albers CA, Banks E, DePristo MA, Handsaker RE, Lunter G, Marth GT, Sherry ST, et al: The variant call format and VCFtools. Bioinformatics 2011, 27:2156–2158.

43. Purcell S, Neale B, Todd-Brown K, Thomas L, Ferreira MA, Bender D, Maller J, Sklar P, de Bakker PI, Daly MJ, Sham PC: PLINK: A tool set for whole-genome association and population-based linkage analyses. Am J Hum Genet 2007, 81:559–575.

44. Alexander DH, Novembre J, Lange K: Fast model-based estimation of ancestry in unrelated individuals. Genome Res 2009, 19:1655–1664.

45. Durbin RM, Abecasis GR, Altshuler DL, Auton A, Brooks LD, Gibbs RA, Hurles ME, McVean GA: A map of human genome variation from population-scale sequencing. Nature 2010, 467:1061–1073.

46. Paschou P, Ziv E, Burchard EG, Choudhry S, Rodriguez-Cintron W, Mahoney MW, Drineas P: PCA-correlated SNPs for structure identification in worldwide human populations. PLoS Genet 2007, 3:1672–1686.

47. Elhaik E: In search of the jtidische Typus: a proposed benchmark to test the genetic basis of Jewishness challenges notions of “Jewish biomarkers”. Front Genet 2016, 7.

48. Marshall S, Das R, Pirooznia M, Elhaik E: Reconstructing Druze population history. Scientific Reports 2016, 6:35837.

49. Das R, Wexler P, Pirooznia M, Elhaik E: Localizing Ashkenazic Jews to primeval villages in the ancient Iranian lands of Ashkenaz. Genome Biol Evol 2016, 8:1132–1149.

50. Marcus JH, Novembre J: Visualizing the geography of genetic variants. Bioinformatics 2017, 33:594–595.

